# Validating metabarcoding-based biodiversity assessments with multi-species occupancy models: a case study using coastal marine eDNA

**DOI:** 10.1101/797852

**Authors:** Beverly McClenaghan, Zacchaeus G. Compson, Mehrdad Hajibabaei

**Affiliations:** Centre for Environmental Genomics Applications, eDNAtec Inc., St. John’s, NL, Canada; Centre for Biodiversity Genomics & Department of Integrative Biology, University of Guelph, Guelph, ON, Canada

**Keywords:** environmental DNA, occupancy modelling, DNA metabarcoding, model selection, marine biomonitoring

## Abstract

Environmental DNA (eDNA) metabarcoding is an increasingly popular method for rapid biodiversity assessment. As with any ecological survey, false negatives can arise during sampling and, if unaccounted for, lead to biased results and potentially misdiagnosed environmental assessments. We developed a multi-scale, multi-species occupancy model for the analysis of community biodiversity data resulting from eDNA metabarcoding; this model accounts for imperfect detection and additional sources of environmental and experimental variation. We present methods for model assessment and model comparison and demonstrate how these tools improve the inferential power of eDNA metabarcoding data using a case study in a coastal, marine environment. Using occupancy models to account for factors often overlooked in the analysis of eDNA metabarcoding data will dramatically improve ecological inference, sampling design, and methodologies, empowering practitioners with an approach to wield the high-resolution biodiversity data of next-generation sequencing platforms.

## Introduction

Environmental DNA (eDNA) as a signal for diversity detection is rapidly advancing. In freshwater systems, in particular, eDNA is now used as a bioassessment tool in both single-species qPCR-based studies and in sequencing-based metabarcoding community assessments [1–3]. Approaches based on eDNA are also gaining traction in the marine environment [4,5]. Oceans are complex, highly diverse, and difficult to sample; therefore, identifying organisms from all trophic levels and taxonomic groups from a single survey method will greatly facilitate rapid, consistent biodiversity surveys [6]. eDNA metabarcoding provides a streamlined method of biodiversity assessment, generating high-resolution biodiversity data with time and effort savings during sample collection and analysis [7,8].

However, there are several levels of uncertainty associated with eDNA sampling for community assessments. The potential for false negatives during sampling, where a species present in the environment is not detected in surveys, can bias results [9]. False negatives can occur during field sampling and during lab processing. If imperfect detection is not accounted for, this could lead to biased estimates of species richness and individual species occupancy [10,11]. Accounting for false negatives will improve community-wide species occurrence estimates based on eDNA surveys and yield more robust ecological conclusions for making management decisions and informing sampling designs. Optimal sampling designs for eDNA metabarcoding studies are not well-established and differ from traditional ecological sampling methods in the cost and effort required for sample collection [12]. Additionally, there are several added variables that need to be accounted for in metabarcoding studies compared to traditional sampling approaches, such as sequencing depth and marker selection, which vary between studies and can affect metabarcoding results [5,13,14]. Sampling designs should be experimentally informed and optimized specifically for eDNA metabarcoding methods [15], yet this is seldom practiced, and these added sources of variation during sample processing are seldom considered in the same analysis as sampling design.

Occupancy modelling is a powerful tool to account for the additional sources of variation associated with next-generation biomonitoring approaches, and it has been used to assess imperfect detection in terrestrial bioassessment [16–18]. These models include 2-levels: the probability that a species occurs at a site (occupancy; ψ) and the probability of detecting a species at a site (probability of detection; *p*). Recently, occupancy models have been adapted for single-species eDNA studies, where occupancy refers to the probability of a species’ DNA occurring at a site, probability of detection refers to the probability of detecting a species in a PCR replicate, and an additional stochastic level is added to assess the probability of capturing a species’ eDNA in a field sample (probability of capture, ϴ; [19,20]) The use of occupancy models in single-species eDNA studies is not ubiquitous, but it is increasing [21].

Occupancy modelling can also be applied to whole communities through multi-species occupancy models, which are commonly applied to traditional surveys in terrestrial systems [22,23], yet seldom used in the context of DNA metabarcoding (Supporting Information 1). In the same way that single-species models were adapted for eDNA studies through the inclusion of an additional stochastic level, multi-species models can be adapted for metabarcoding by including this additional level. Modeling communities together in a single multi-species model can improve the accuracy and predictive ability of occupancy models compared to single-species models [24]. Application of multi-species, multi-scale occupancy models to metabarcoding data are rare, focusing on small-scale lab manipulations [25], and no studies have implemented this modelling approach to improve sampling designs in natural systems (but see [26] for a single species example). Incorporating these models routinely in metabarcoding analysis will improve ecological inferences and species richness estimates, as well as facilitate the development of robust sampling designs for a relatively new technique where little thought has been dedicated to developing de novo sampling methods distinct from traditional sampling methods. The inclusion of covariates in occupancy models at each process level extends the application of the model, enabling discrimination between sources of variation in sampling effort and environmental factors. However, making conclusions based on models with covariates requires methods of model assessment and selection for multi-species, multi-scale models.

Here, we demonstrate how multi-species occupancy modelling can be used for the analysis of community biodiversity data resulting from eDNA metabarcoding and highlight the potential of these models for both improving methodologies and sound ecological inference. We present methods for model assessment and model comparison adapted for multi-scale, multi-species occupancy models. Finally, we demonstrate how these tools can improve inferential power from eDNA metabarcoding results using a case study in a coastal, marine environment.

## Material & Methods

### Model Formulation

#### The multi-species, multi-scale occupancy model

We used a Bayesian modeling framework to develop a multi-species, hierarchical occupancy model with three stochastic levels: occupancy (*ψ*), probability of capture (*ϴ*), and probability of detection (*p*) (Figure 1). The occupancy process describes whether sampling sites are occupied or not by a given species’ DNA. For eDNA sampling, there are often two levels of sampling replication within each site (e.g. [20,27]): biological replicates are samples collected from a single site in the field and technical replicates are repeated samples taken from a single biological replicate in the lab. The probability of capture refers to the probability that a species’ DNA is collected in a sample, given that the species was present at the site. The probability of detection refers to the probability that a species was detected in a technical replicate, given that the species’ DNA was collected in the sample. This model assumes no false positives occur in the data. While false positives may be a possibility in metabarcoding data [15], we used strict bioinformatic filtering to reduce this possibility (see *Bioinformatics* below). Further comments on false positives can be found in the *Discussion*.

**Figure 1.**
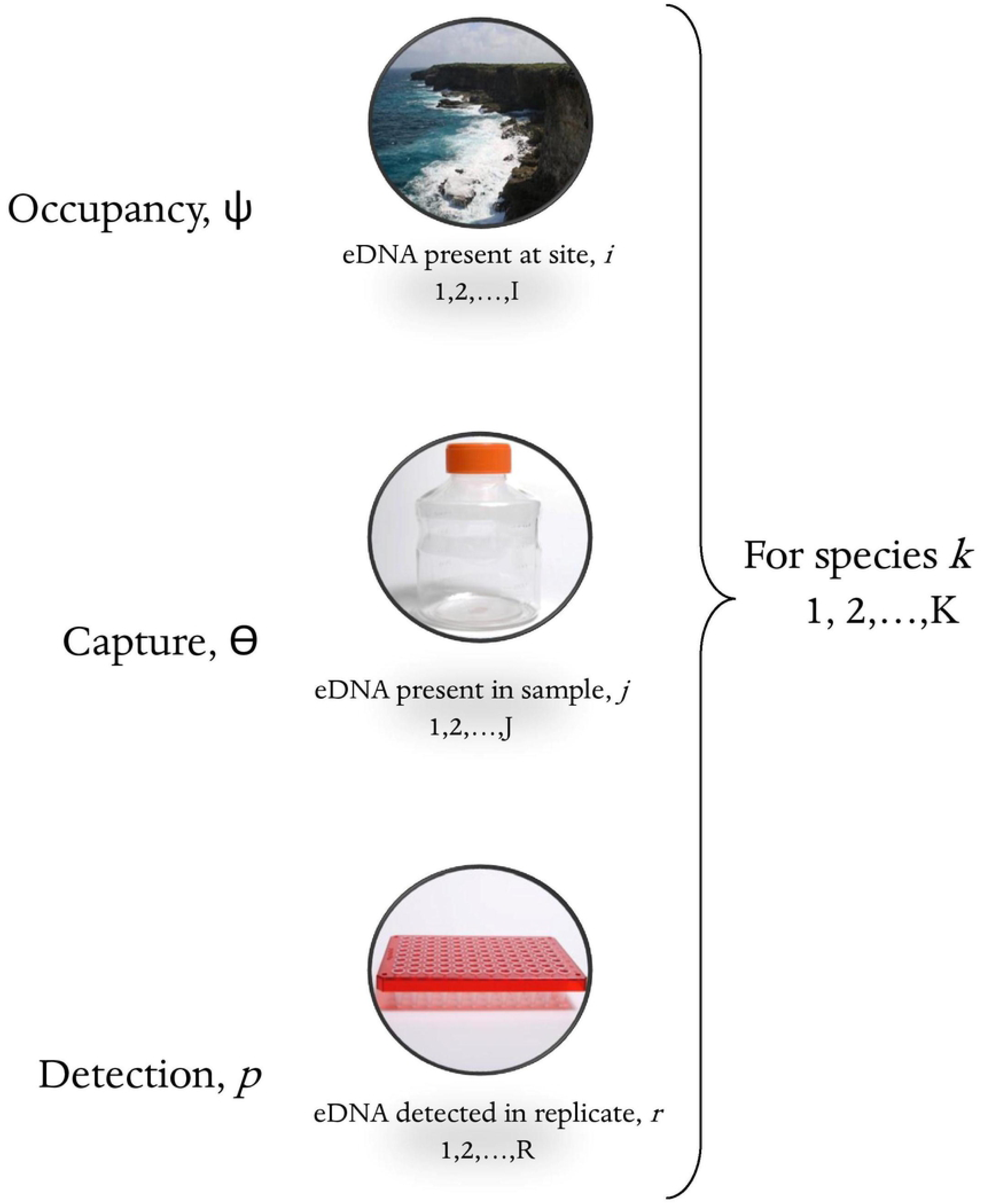
Schematic illustration of the three stochastic levels included in the multi-scale, multi-species occupancy model.

This model can be fit to a dataset, *y*_*ijrk*_, which is a binary indicator of whether a species *k* (*k* = 1,2,…K) was detected (1) or not detected (0) in a technical replicate *r* (*r* = 1,2,…R) from a given sample *j* (*j* = 1,2,…J) at a given site *i* (*i* = 1,2,…I). The model consists of three coupled Bernoulli trials to describe a four-dimensional array of data *y*_*ijrk*_.

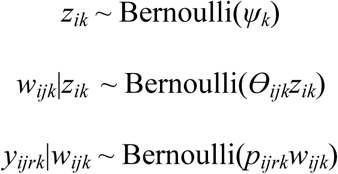

The first random variable *z*_*ik*_ describes the detection (*z*_*ik*_ = 1) or non-detection (*z*_*ik*_ = 0) of species *k* at site *i* as a function of the occupancy probability *ψ*_*k*_. The second random variable *w*_*ijk*_ describes the detection (*w*_*ijk*_ = 1) or non-detection (*w*_*ijk*_ = 0) of species *k* in sample *j* at site *i* as a function of the probability of capture (*ϴ*_*ijk*_) and the occupancy state (*z*_*ik*_).

Covariates can be included in the model at each stochastic level (e.g., α1, α2, α3). Continuous covariates were z-score standardized to have a mean of zero and a standard deviation of one to help with model convergence. Categorical covariates can also be included at any level, which is demonstrated below at the probability of detection level (i.e., α4). Covariates are included in the model as follows:

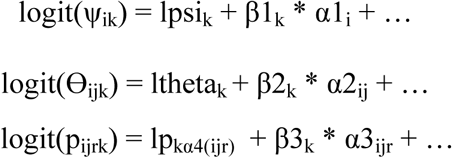

For multi-species occupancy models, species coefficients arise from additional community-level parameters:

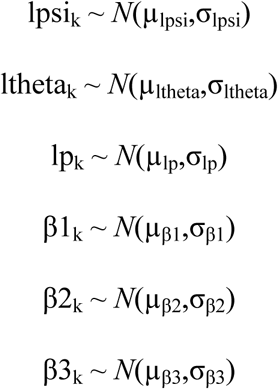

Community-level parameters are described by weakly informative hyperpriors [28]. All mean values for the above prior distributions were selected from a normal distribution and all standard deviations were selected from a uniform distribution.

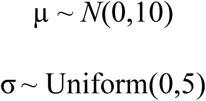

Prior sensitivity was assessed by running the model with various prior parameterizations. Posterior distributions were similar across all priors.

#### Model Assessment and Comparison

To assess model fit, we looked at diagnostic plots to examine model fit and highlight areas of lack of fit. We plotted the deviance residuals for each species and site, and plotted deviance residuals against covariates. We calculated Bayesian *p*-values following [29], adapted for a multi-scale model (Supporting Information 2) to assess goodness-of-fit, where values close to 0.5 indicate a good fit and values >0.95 or <0.05 indicate a poor fit.

We also adapted model selection and cross-validation calculations from [29] for multi-scale, multi-species occupancy models to determine the best model. We calculated the Watanabe-Akaike information criterion (WAIC; [30]) and the conditional predictive ordinate criterion (CPO; [31]), and then evaluated the results of *k*-fold cross validation using the Brier score and the logarithmic score. The complete calculations for all model assessment and comparison methods can be found in Supporting Information 2.

#### Unknown Species Richness

In addition to the model described above, we implemented a model using data augmentation for communities with unknown species richness [10]. This model can be used to estimate species richness for the sampling area through the inclusion of another Bernoulli variable:

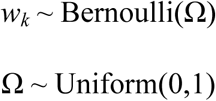

For species *k* (*k* = 1,2,…*M*), *M* is the total number of species in the augmented model and *w*_*k*_ = 1 if species *k* was ever detected during the study. An upper limit to species richness (*M*) is specified a priori and considered large enough when the estimate of true species richness is sufficiently lower than *M* (i.e., the value of *M* is in the right tail of the posterior distribution of species richness; [28]).

### Case Study: Conception Bay, Newfoundland

#### Sample Collection, Processing and Sequencing

Triplicate 250 mL water samples were collected from coastal surface water at eight sites along two transects in Conception Bay, Newfoundland and Labrador, Canada, on October 13–14, 2017. Water samples were filtered using 0.22 μm PVDF Sterivex filters (MilliporeSigma) and DNA was extracted from filter membranes using the DNeasy PowerWater Kit (Qiagen). Five target markers in the cytochrome *c* oxidase I (COI) region were amplified by PCR from each sample. Table 1 details the primer sets used to target these markers. Three PCR replicates were performed for each amplicon from each sample and then pooled for a single PCR cleanup with the QIAquick 96 PCR purification kit (Qiagen). Amplicons were then indexed using unique dual Nextera indexes (IDT). All amplicons were pooled into one library to normalize DNA concentration and the library was sequenced with a 300-cycle S4 kit on the NovaSeq 6000 following the NovaSeq XP workflow. Raw sequence reads are available in NCBI’s sequence read archive under accession number PRJNA574050. Primers were trimmed from sequences and then DADA2 v1.8.015 [32] was used for quality filtering, joining paired end reads and denoising to produce exact sequence variants (ESVs). Taxonomy was assigned using NCBI’s blastn tool v2.6.026 [33] to compare ESV sequences against the nt database. See [5] for detailed sampling, sequencing, and bioinformatic methodology.

**Table 1.**
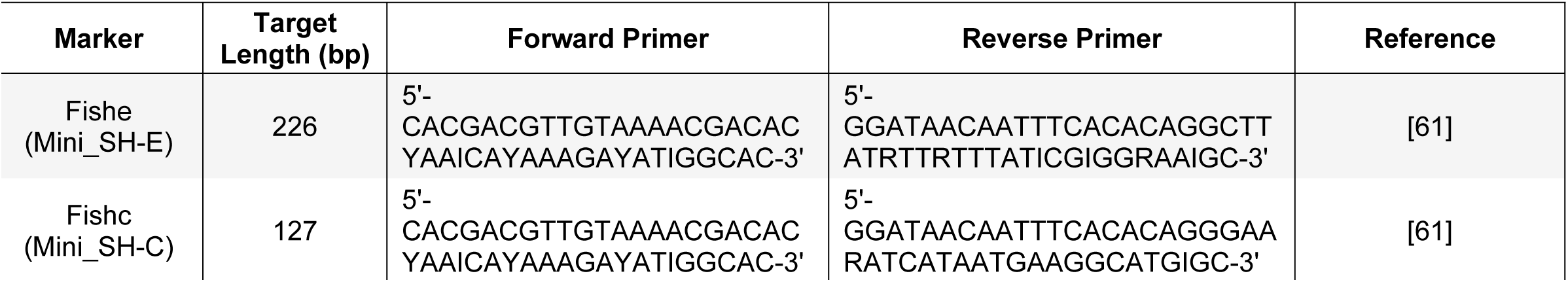
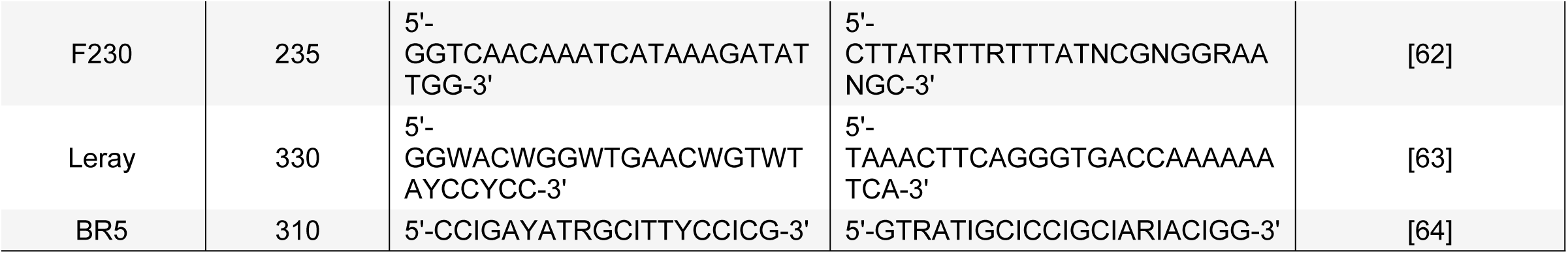
Primer pairs used to amplify five target amplicons in the COI region of the mitochondrial genome from water samples collected in Conception Bay, Newfoundland, Canada.

#### Occupancy Model Implementation

Under the occupancy modelling framework described above, each collection site along each transect in Conception Bay was considered a different site in the occupancy model. Replicate bottles collected at a site were considered samples. Each amplicon sequenced from each bottle was considered a technical replicate. While we conducted replicate PCRs of each amplicon, the products were pooled prior to sequencing so we did not include PCR replicates separately in our models. However, PCR replicates can easily be accommodated in multi-scale, multi-species occupancy models, such as the model described here.

We included sequencing depth (number of reads per sample per amplicon) as a continuous covariate at the level of probability of detection. Additionally, we included amplicon identity as a categorical covariate at the level of probability of detection. We included water depth (m) as a continuous covariate at the level of occupancy. We compared a null model with no covariates with four models with different combinations of covariates (Table 2).

**Table 2.** Model comparison between multi-scale, multi-species occupancy models using four methods (WAIC, CPO, Brier Score and Log Score). The covariates (water depth at the sampling site, sequencing depth for each technical replicate, and amplicon sequenced for each technical replicate) included at each level of the model (occupancy: ψ, capture: ϴ, detection: p) are listed on the left. Bolded values indicate the best model for each method of model comparison.

| MODELS | WAIC | CPO | Brier Score | Log Score |
| --- | --- | --- | --- | --- |
| **Model 1**<br>$\psi(.) \Theta(.) p(.)$ | 16633 | 2904627 | 293 | 2291 |
| **Model 2**<br>$\psi(\text{water depth}) \Theta(.) p(\text{sequencing depth, amplicon})$ | 62255 | 8069266 | 334 | 3715 |
| **Model 3**<br>$\psi(.) \Theta(.) p(\text{sequencing depth})$ | **16184** | 2395664 | 291 | 2279 |
| **Model 4**<br>$\psi(.) \Theta(.) p(\text{amplicon})$ | 61864 | 9310577 | 333 | 3842 |
| **Model 5**<br>$\psi(\text{water depth}) \Theta(.) p(.)$ | 16348 | **2027311** | **283** | **2188** |

All statistical analyses were conducted in R v3.5.1 [34]. MCMC sampling was achieved with JAGS [35], implemented using ‘*jagsUI’* v1.5.0 [36]. The model was written for JAGS in the BUGS language (see Supporting Information 3 for BUGS model structure of the most complex model). We fit models using known species richness to conduct our model comparisons, and assessed models and model fit to determine the best model. MCMC sampling was run in three chains, each with 50,000 iterations, a burn in of 10,000, and a thinning rate of 10. Convergence was verified using the Gelman-Rubin diagnostic [37] and by evaluating trace plots. For all models, we report parameter estimates as the mean of the posterior distribution with the 95% highest posterior density interval (HDI; [38]) calculated using ‘HDInterval’ v0.2.0 [39]. Significance of continuous covariates was assessed by determining if the 95% confidence intervals of parameter estimates overlapped with zero [28]. For the categorical covariate amplicon, we used a generalized linear model with a beta distribution implemented using ‘*betareg’* [40] to compare the estimated species-specific probabilities of detection between markers and phyla. Likelihood ratio tests were used to determine the significance of predictors at α = 0.05. We conducted a data augmented model with unknown species richness for the best model at varying levels of augmentation to determine the minimal level of augmentation required, as described above in the *Unknown Species Richness* section.

## Results

We ran five multi-species, multi-scale occupancy models with different combinations of covariates (i.e., water depth at the level of occupancy, sequencing depth and amplicon at the level of detection probability) and assessed these models using model comparison and cross-validation methods adapted for this multi-scale approach (Table 2). Three of the model comparison methods (CPO and two cross-validation scores) were in agreement that Model 5 (*ψ(water depth) ϴ(.) p(.)*) was the best model, while the WAIC suggested Model 3 (*ψ(.) ϴ(.) p(sequencing depth)*) was the best model. We considered Model 5 our best model moving forward, given that most selection methods indicated this was the best model.

We assessed model fit using Bayesian *p*-values and diagnostic plots for all models but present the results for the best model only. We obtained a Bayesian *p*-value of 0.51, suggesting that Model 5 (*ψ(water depth) ϴ(.) p(.)*) provided a good fit to our data overall; diagnostic plots, however, revealed higher deviance at sites with lower water depth, suggesting a poorer model fit at shallower sites (Supporting Information 4). The community-wide estimate for occupancy was 0.27 (HDI: 0.22-0.33). Water depth had a significant effect on the community mean occupancy (Figure 2), and we detected considerably more species at the shallowest sites compared to the other sites (274 species at two shallow water sites combined compared to 109 species across all six deep water sites). The community-wide probability of capture was 0.98 (HDI: 0.96-0.99) and the community-wide probability of detection was 0.15 (HDI: 0.14-0.17). Species-specific estimates of occupancy, capture probability, and detection probability were also obtained from the model (Supporting Information 5).

**Figure 2.**
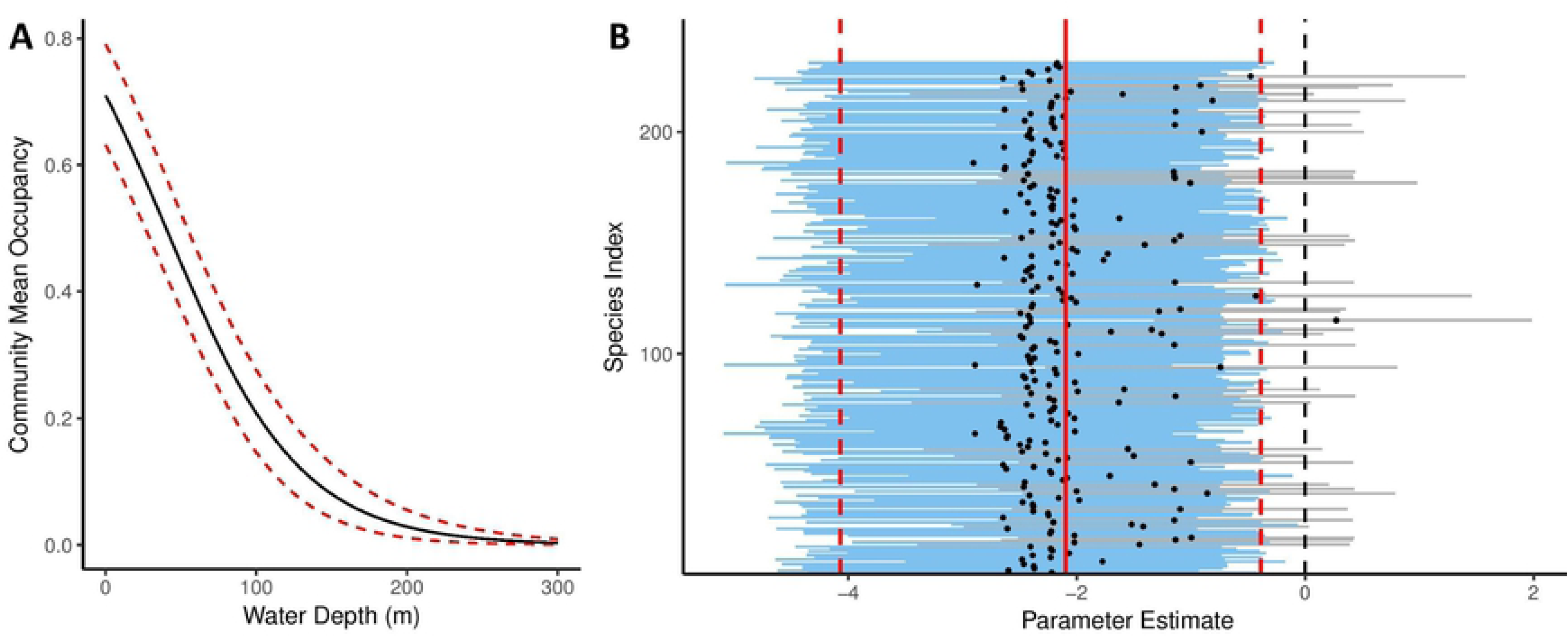
(A) Community mean occupancy by water depth (m) predicted using a multi-species, multi-scale community occupancy model. The gray area represents the 95% confidence interval. (B) Parameter estimate for each species for the effect of water depth on occupancy in a multi-species, multi-scale community occupancy model. Solid red line indicates the community mean and dashed red lines indicate the upper and lower limits of the 95% confidence intervals of the community mean parameter estimate. Blue lines indicate 95% confidence intervals of individual species parameter estimates that do not overlap with 0. Grey lines indicate 95% confidence intervals of individual species parameter estimates that do overlap with 0.

While it was not selected as our best model, we present the results from Model 4 (*ψ*(.) *ϴ*(.) *p*(*amplicon*)) to demonstrate how categorical covariates can be incorporated into the occupancy modelling framework. Amplicons displayed significantly different probabilities of detection (*X*^2^ = 34.43, p-value < 0.001; Figure 3). When considering species-specific probabilities of detection and including phylum-level identifications, there was a significant interaction between amplicon and phylum (*X*^2^ = 85.18, p-value < 0.001), and some amplicons clearly failed to detect certain taxonomic groups (Figure 4).

**Figure 3.**
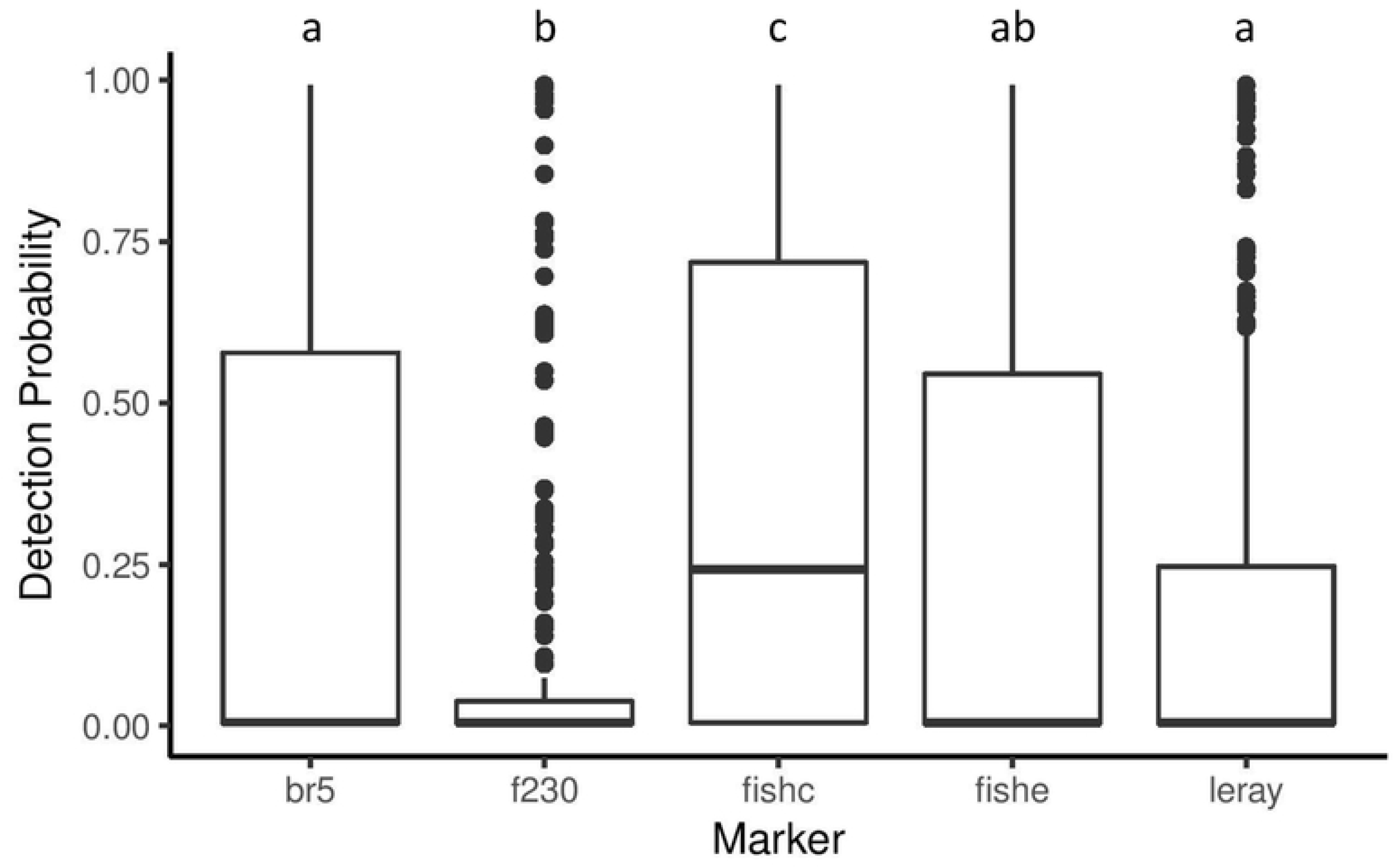
Mean detection probability estimated from occupancy model 3 (ψ(.) ϴ(.) p(amplicon)) for each species plotted by amplicon. The band in the middle of the box represents the median and the upper and lower edges of the box represent the upper and lower quartiles. The whiskers represent 1.5 times the inter-quartile range. Beta regression indicated a significant effect of amplicon on probability of detection (X^2^ = 34.43, p-value < 0.001). Significant different (α = 0.05) between amplicon are denoted by different letters above each amplicon.

**Figure 4.**
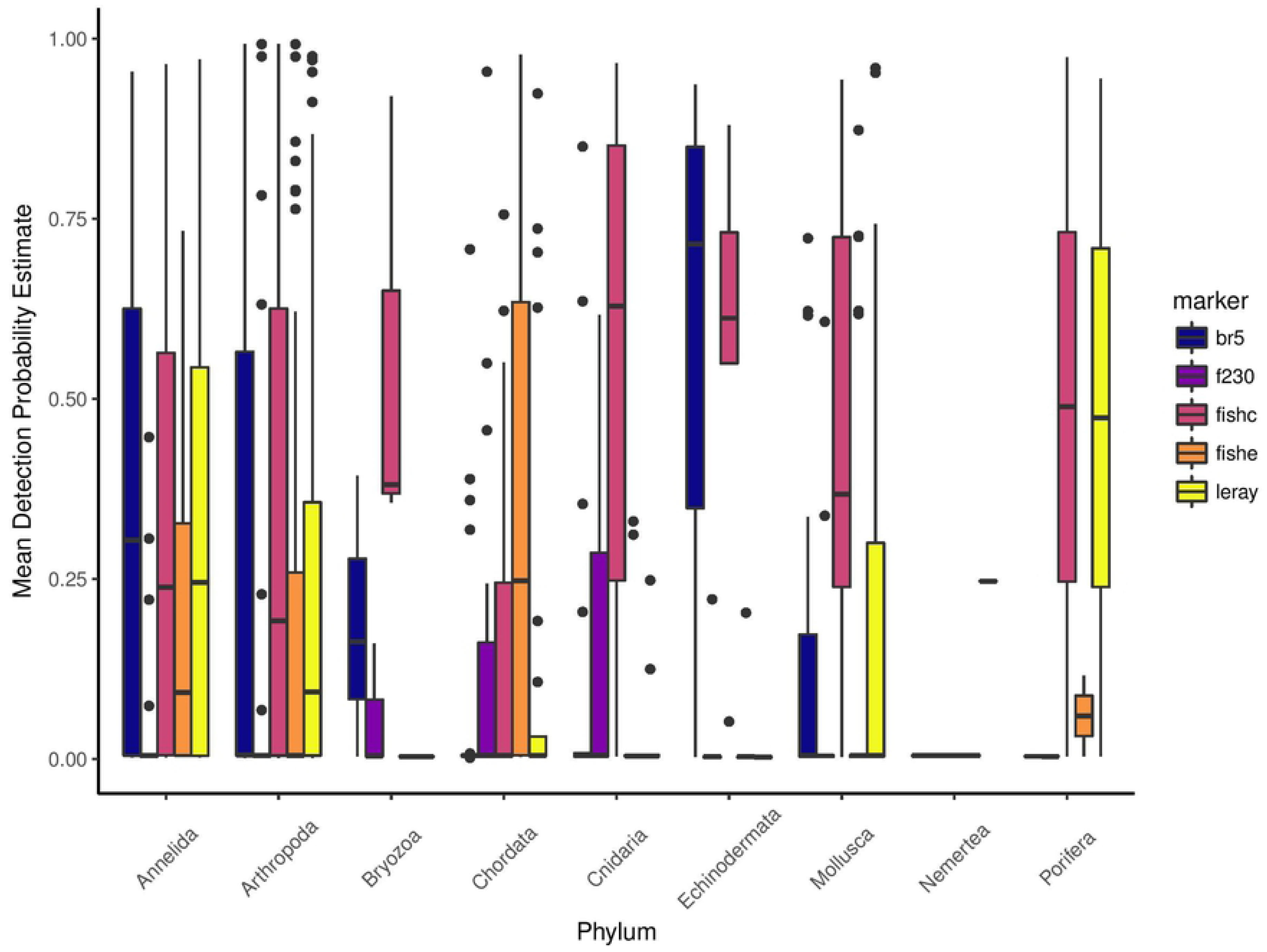
Mean detection probability for each species plotted by amplicon and phylum for metazoan phyla only. The band in the middle of the box represents the median and the upper and lower edges of the box represent the upper and lower quartiles. The whiskers represent 1.5 times the inter-quartile range.

Sequencing depth was not included as a covariate in the best model; in the best model that did include sequencing depth, Model 3 (*ψ(.) ϴ(.) p(sequencing depth)*), we observed no significant effect of sequencing depth in this case study (Supporting Information 6).

We estimated species richness for the survey area by running the best model with data augmentation. This model used the probabilities of capture and detection to estimate the number of species missed in sampling efforts. We detected 231 species overall, and the estimated species richness for the survey area was 284 (HDI: 262-307), indicating that 53 (HDI: 31-76) species were undetected during our surveys. In other words, our survey detected ~81% of the estimated species in our study area.

## Discussion

We applied a multi-species, multi-scale occupancy model to a DNA metabarcoding dataset generated from marine water samples and explored how the inclusion of categorial and continuous covariates at different levels improved model performance. The best model included water depth as a covariate at the level of occupancy, where we observed a higher species richness at shallower sites. One of the shallow water collection sites was within 1 km of a sewage outflow, which may have contributed to this result, although a high species richness was also observed at the second, shallow water site located >10 km from the sewage outflow. The probability of capture estimate of 0.98 suggests a high probability of collecting a species’ DNA in a given sample. However, the detection probability was relatively low at 0.15, likely because many species were not detected consistently by multiple amplicons, and a low probability of detection can lead to overestimates for higher level parameters [41].

We observed a significant effect of amplicon and phylum on the species-specific probabilities of detection. Since the performance of each amplicon varies by taxonomic group (this study; [13]), including a variety of target regions is important to detect species across the tree of life, and increasing the number of technical replicates using a target region will not necessarily improve the community-wide probability of detection. We observed no significant effect of sequencing depth in this study. However, the samples were all sequenced on a NovaSeq instrument, which generates an unprecedented number of reads, yielding very high sequencing depths (mean number of filtered sequences per sample ± standard deviation: 8,519,055 ± 2,514,998) compared to many other barcoding studies (e.g. [42,43]). In studies where the mean sequencing depth is lower, differences in sequencing depth are likely to have greater effects [5,44].

We used the occupancy modeling framework to estimate the species richness for the survey area and determined that 53 species or approximately 19% of the estimated number of species present were undetected during our surveys. Similar to many ecological studies, the case study presented here included a relatively low spatial coverage (*n* = 8 sites), but our occupancy modelling approach allowed us to assess false absences in our study, which is a significant improvement from most metabarcoding surveys [11]. The proportion of species detected could be improved by (1) increasing sampling effort in the field by sampling more sites, (2) collecting more replicate biological samples at each site, and (3) including additional target regions during laboratory processing. Given the limited extent and breadth of our sampling effort, the conclusions regarding the effect of covariates and the estimates of occupancy, capture, and detection probabilities for individual species should not be extrapolated to other systems. Further research should investigate the impacts of variation in sequencing depth and target regions on detection probability in metabarcoding studies, particularly in other ecosystems and across greater spatial scales.

Through the inclusion of environmental and experimental covariates, the multi-species occupancy framework can be applied for direct ecological assessment and to improve the methodology for next-generation biodiversity assessment. From an ecological perspective, environmental variables (e.g. temperature, salinity, turbidity) can be included at the level of occupancy to determine their effects on community diversity and the presence of individual species. From a methodological perspective, environmental and experimental variables (e.g. sample volume, sequencing depth) can be included at the level of field sampling and technical replication to understand how these factors affect metabarcoding results. Understanding the effects of these covariates facilitates the development of more robust experimental and survey designs. Furthermore, simulations using occupancy models can be used to optimize sampling effort, enabling practitioners to fine-tune the trade-off between field sampling and lab work [21]. The number of sites, biological samples, and technical replicates can all be optimized to maximize the species richness recovered from eDNA samples. PCR level stochasticity, which is known to affect sequencing results [44,45], was not considered in our case study (i.e., PCR replicates were pooled before sequencing) but PCR replicates can easily be included as technical replicates in the model described here. PCR replicates are commonly included separately in single-species occupancy models for eDNA data [19,20,27]. By including PCR replicates as technical replicates, additional stochasticity in the sampling process can be accounted for, further improving inferences.

A key advantage of the occupancy modeling framework demonstrated here is its flexibility. Modifications to the model can allow several additional factors to be included, and a priori information can be used to guide model development. For example, multiple sampling periods have been included in dynamic, multi-season occupancy models to quantify temporal changes in community structure (e.g. [22]). Repeated eDNA sampling for metabarcoding could be modelled similarly to account for local extinction and colonization events between sampling periods. In addition to accounting for false negatives, several studies have developed methods for including false positives in occupancy models [46–48]. False positives may potentially arise from metabarcoding data through sequencing errors, PCR errors, and poor reference database coverage or quality [15,49,50]. Strict bioinformatic filtering helps to minimize the inclusion of these errors in resulting data sets; however, the possibility of false positives cannot be eliminated. Our model did not consider false positives, and, to our knowledge, these have yet to be incorporated into multi-species occupancy models. The occupancy modeling framework can also be adapted to include or estimate taxa abundances [28]. Following current protocols, abundance estimates from metabarcoding data are not reliable [51,52], but these models may provide tools to improve abundance estimates from metabarcoding data.

We demonstrate for the first time how a multi-scale, multi-species occupancy modelling framework can be used in a natural system to account for imperfect detection and allow for critical assessment of experimental and environmental factors influencing biodiversity data from eDNA metabarcoding. Despite the utility of these models for improving detection and targeting areas of variation in the pipeline from sample collection to sample processing, this approach has been underutilized in DNA metabarcoding studies (Supplementary Information 1; but see [25]). This multi-species occupancy modelling framework will be particularly useful for bioassessment studies using DNA metabarcoding because it will improve estimates of occupancy and species richness, aid in optimizing sampling efforts in the field and lab, and, using the model assessment methods described here, identify ecological and environmental factors affecting occupancy, capture, and detection probabilities. Given the high stakes for documenting and understanding biodiversity that is under increasing anthropogenic threat [53] and decline [54] globally, new tools are imperative for rapid bioassessment [7,55,56]; yet, like any emergent technology, there is the potential to misuse these tools [57], which can have unforeseen consequences (e.g. [58]). In the case of DNA metabarcoding, neglecting to assess imperfect detection at key points along the sample collection and processing pipeline could lead to failure to detect species of interest, biased estimates of species richness, and miscalculations of species distributions, all of which have consequences for conservation and management [24,59,60]. We recommend incorporating multi-scale, multi-species occupancy modeling into the design and analysis of future metabarcoding studies.

## Acknowledgements

We would like to thank Nicole Fahner, Joshua Barnes, Avery McCarthy, Greg Singer, Hoda Rajabi, and Emily Porter for their assistance in sample collection, processing and bioinformatics.

